# A split GFP system to enhance spatial and temporal sensitivity of translating ribosome affinity purification (TRAP)

**DOI:** 10.1101/2021.01.29.428889

**Authors:** Kasia Dinkeloo, Zoe Pelly, John M. McDowell, Guillaume Pilot

## Abstract

Translating ribosome affinity purification (TRAP) utilizes transgenic plants expressing a ribosomal protein fused to a tag for affinity purification of ribosomes and the mRNAs that they are translating. These actively translated mRNAs (translatome) can be interrogated by qPCR or RNAseq. Condition- or cell-specific promoters can be utilized to isolate the translatome of specific cell types, at different growth stages and/or in response to environmental variables. While advantageous for revealing differential expression, this approach may not provide sufficient sensitivity when activity of the condition/cell-specific promoter is weak, when ribosome turnover is low in the cells of interest, or when the targeted cells are ephemeral. In these situations, expressing tagged ribosomes under the control of these specific promoters may not yield sufficient polysomes for downstream analysis. Here, we describe a new TRAP system that employs two transgenes: one is constitutively expressed and encodes a ribosomal protein fused to one fragment of a split GFP; the second is controlled by a stimulus-specific promoter and encodes the second GFP fragment fused to an affinity purification tag. In cells where both transgenes are active, the purification tag is attached to ribosomes by bi-molecular folding and assembly of the split GFP fragments. This approach provides increased sensitivity and better temporal resolution because it labels pre-existing ribosomes and does not depend on rapid ribosome turnover. We describe the optimization and key parameters of this system, and then apply it to a plant-pathogen interaction in which spatial and temporal resolution are difficult to achieve with current technologies.

**Significance:** Translating ribosome affinity purification (TRAP) has been modified to allow with increased sensitivity the isolation of RNA from sets of cells in which the activity of condition/cell-specific promoters is weak, ribosome turnover is low, or cells whose nature is ephemeral. Based on the use of a split linker constituted of the GFP driven by a pathogen-inducible promoter, this new TRAP system enabled efficient isolation of translated RNA from pathogen-infected leaf cells.

## Introduction

<u>T</u>ranslating <u>r</u>ibosome <u>a</u>ffinity <u>p</u>urification (TRAP) was pioneered in mice (Heiman, Schaefer et al. 2008) and adapted for plants as a technique to reproducibly isolate actively-translated mRNAs from genetically defined populations of cells (Kage, Powell et al. 2020). TRAP utilizes transgenic organisms expressing a tagged ribosomal protein that is incorporated into ribosomes by the cell. Cell constituents are fractionated under conditions that preserve associations between ribosomes and the associated mRNAs (*i.e.* polysomes). Tagged polysomes are then purified by immunoprecipitation using an antibody against the tag. RNA extracted from the immunoprecipitated polyribosomes contains the translatome, *i.e.* the collection of mRNAs that are associated with and being translated by ribosomes (sometimes referred to as ribosome-nascent chain complex bound mRNAs, or RNC-mRNAs). Conventional transcriptomics captures all mRNAs, including those that are not mature or actively translated at the time of isolation. Thus, traditional transcriptomics cannot capture the control that an organism exerts on translation (Adams 2008). Contrastingly, the translatome correlates more closely to the proteome (Wang, Cui et al. 2013) and can therefore provide a more accurate assessment of physiological status. In addition, conventional transcriptomic analyses are typically performed using mRNA purified from homogenized organs comprised of various differentiated cell types. With this approach, it is impossible to distinguish from one another the transcriptomes of the various cell types that comprise the organ. On the other hand, TRAP can be customized to reveal the translatome in specific cell types by expressing the tagged ribosomal protein under the control of a cell type-specific promoter. Similarly, responses to environmental stimuli can be assessed by expressing the ribosomal protein using a stimulus-dependent promoter. In these approaches, the organ is harvested and the tagged polysomes are purified from the mixed population of tagged and untagged polysomes (Zanetti, Chang et al. 2005, Mustroph, Juntawong et al. 2009) (Figure 1a). This approach enriches for polysomes from the cell type of interest because the tagged ribosomal protein is expressed only in the specific cell type(s) or conditions in which the promoter is active. mRNA is then isolated from the immunopurified polysomes and subjected to RNA sequencing (or other analyses), yielding cell type- or stimulus-specific translatomes.

**Figure 1.**
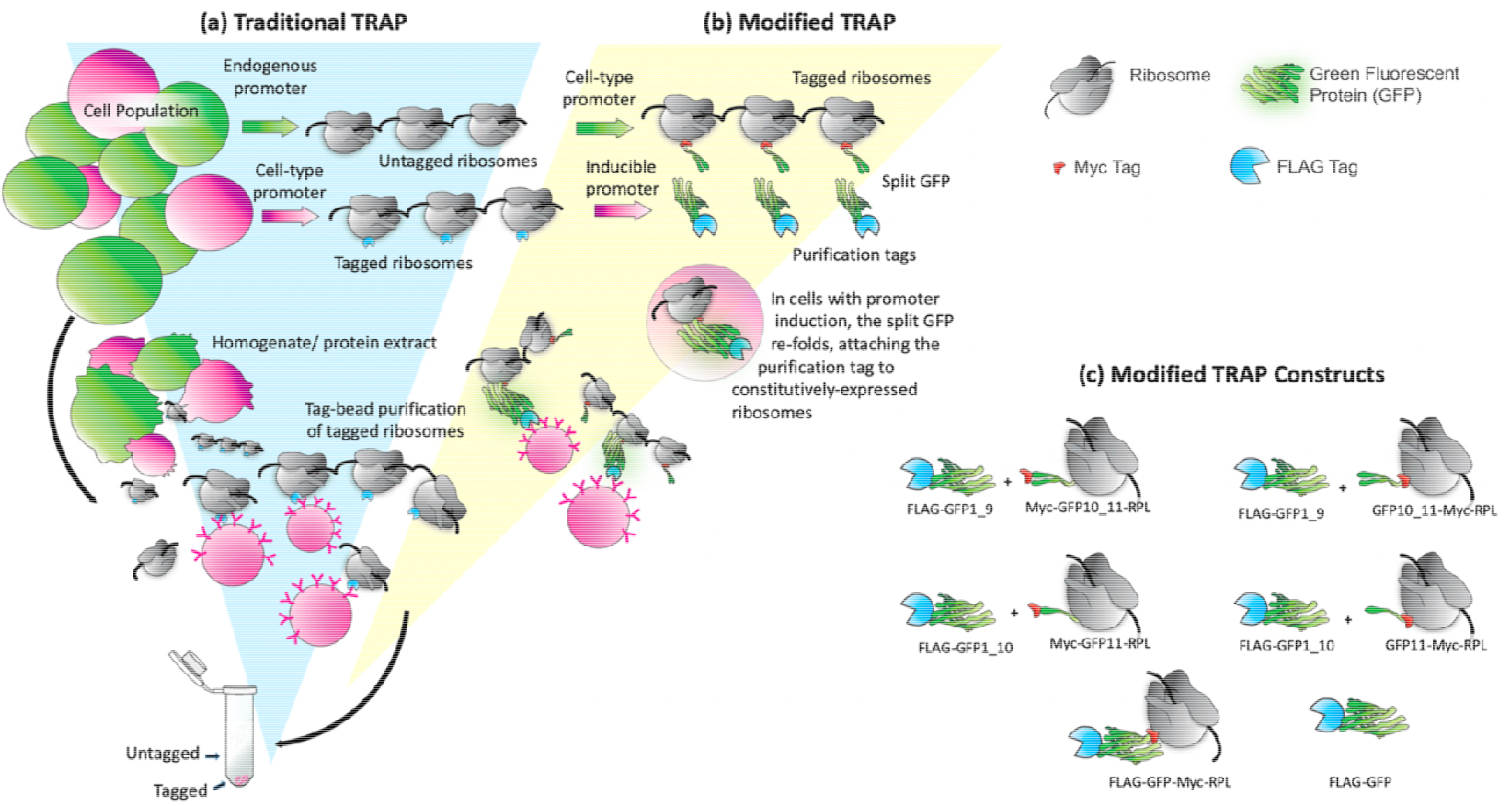
Schematic of traditional and split GFP TRAP. (a) In traditional TRAP, a promoter drives expression of a ribosomal protein fused to an epitope tag. The tag allows for affinity purification of polyribosomes and associated RNAs. Placing the tagged ribosomal protein gene under the control of a conditionally active promoter (red) can provide enrichment of ribosomes from specific cells of interest (red) in a complex mixture (green) (b) The split GFP TRAP system employs two transgenes: One gene contains a condition-specific promoter that drives the expression of the purification tag and part of a split GFP linker protein (red). The second gene is driven by a constitutive promoter and encodes the remaining portion of the split GFP, fused to an epitope tag and the ribosomal protein for incorporation into a translating ribosome (green). Assembly of the split GFP attaches the purification tag to pre-existing ribosomes, thereby increasing the proportion of tagged ribosomes in cells of interest, compared to the traditional one-gene system that requires replacement of endogenous, untagged ribosomes with those that incorporate the tagged protein. (c) Diagrams representing the constructs used for optimization via transient expression in *N. benthamiana*. In these experiments (Figures 2-5), different splits of GFP and different positions of a constitutive Myc tag were compared.

Applications of TRAP to many organisms, including plants, have shown that translatome analysis can reveal information that is not present in the transcriptome. For example, studies of the translatome in Arabidopsis seedlings exposed to a period of hypoxia identified a specific group of mRNAs that allows for acclimation to hypoxia (Mustroph, Zanetti et al. 2009). These mRNAs were enriched in several specific cell types and encode proteins aiding in stress tolerance, development, and metabolism (Mustroph, Zanetti et al. 2009). TRAP analysis of plant-bacteria interactions demonstrated that translational control is a significant component of immune response regulation and delineated a link between metabolism and plant immune response via specific changes in translation (Meteignier, El Oirdi et al. 2017, Xu, Greene et al. 2017, Yoo, Greene et al. 2020). These studies suggest that additional comparisons of transcriptome and translatome induced by biotic or abiotic stresses will reveal new information that would not be evident with transcriptomic data alone.

Because ribosome biogenesis in plants is not completely understood and ribosomes appear to be relatively stable with a turnover rate of approximately 6-8 days (Salih, Duncan et al. 2020), it may be disadvantageous to directly express the ribosomal protein-purification tag fusion under the control of stimulus-specific promoters. In particular, slow ribosome turnover might retard tagging of active ribosomes with newly expressed, tagged ribosomal proteins, limiting the utility of this technique for short-term treatments. In addition, application of TRAP could be problematic for cells in which ribosomal protein synthesis is repressed, as appears to be the case in cells under stress (Salih, Duncan et al. 2020). To overcome these limitations, we designed an alternative to conventional TRAP in which pre-existing ribosomes could be tagged soon after application of the stimulus. We optimized this system, validated its efficiency, and applied it to the interaction between Arabidopsis and a filamentous oomycete pathogen

## Results

### Design principles of the new TRAP system

Our need for a modified TRAP system arose from the desire to study plant-pathogen interactions between Arabidopsis and its oomycete pathogen, *Hyaloperonospora arabidopsidis* (*Hpa*) (Herlihy, Nora et al. 2019). Specifically, we wanted to profile plant cells that were in direct contact with pathogen feeding structures called haustoria (Bozkurt and Kamoun 2020). These “haustoriated” cells are difficult to analyze because they are present at relatively low abundance even in heavily infected leaves and cannot be purified away from non-haustoriated leaf cells. Moreover, transcriptome data indicates that *Hpa* infection has a suppressive effect on plant ribosome biogenesis (Wang, Barnaby et al. 2011). We reasoned that the potential limitations posed by ribosomal protein turnover and cell-specificity could be addressed by a split-linker protein system encoded by two transgenes (Figure 1b): One constitutively expressed transgene would encode the ribosomal protein fused to one half of the split linker protein. The second gene would be expressed under the control of a stimulus-specific promoter (*e.g.* a pathogen-inducible promoter) and encode the other half of the split linker, fused to a purification tag for TRAP. The split linker would assemble in cells where both transgenes are expressed, effectively fusing the ribosomal protein to the purification tag, thereby enabling the efficient tagging of pre-existing ribosomes from the cells of interest and enrichment from a whole-leaf extract by TRAP.

### Selection of Linker Systems

The success of the strategy described above is predicated upon on stable and strong association of the ribosomal protein with the inducible purification tag via the split linker protein. Therefore, it was important to select a modular linker system from which two protein fragments can be expressed from separate genes and bind to one-another with high affinity and specificity in plant cells. The ideal linker should also be small (less than 50 kDa), nontoxic for plant cells, and the association must withstand purification steps in chaotropic buffers. Several systems meeting these criteria were available: split ubiquitin, streptavidin-streptag-II, iDimerize, PDZ domain-ligand peptides, and split green fluorescent protein (GFP) (Chalfie 1995, Korndörfer and Skerra 2002, Lee and Zheng 2010). Due to issues with potential disruption of cellular functions by ubiquitin, the need for an additional ligand for iDimerize, the size burden of StrepTactin, and a lack of specificity of PDZ domain peptides, we chose the split GFP as a linker.

GFP has been used in many organisms as a reporter due to its intrinsic fluorescence (Chalfie 1995). The protein is composed of 11 ß-sheets that form a barrel around an internal chromophore. Importantly, fragments of GFP can be expressed from two different genes as a split protein that will spontaneously assemble when those fragments are in close proximity, forming a fully active chromophore (Blakeley, Chapman et al. 2012, Ito, Ozawa et al. 2013). “Superfolder” GFP variants are composed of fragments that associate with high affinity, thus comprising a stable linker that would not disassociate during TRAP (Cabantous and Waldo 2006). Moreover, GFP’s intrinsic fluorescence would provide a visual confirmation that both genes are expressed and that the split GFP halves have assembled in the desired cell type. Therefore, we selected the “superfolder” GFP described in (Pedelacq, Cabantous et al. 2006) that refolds robustly from a denatured state, emits fluorescence proportional to protein abundance, and has been validated for efficient and specific assembly from split configurations. This GFP was used previously for a tripartite split protein that was further mutagenized to improve reconstitution and folding efficiency (Cabantous, Nguyen et al. 2013).

In addition to the linker, careful consideration was given to tags and spacers (short amino acid sequences that provide flexibility between the different parts of our system) for the system. Two tags were required: one for affinity purification, positioned under the control of the inducible promoter, and one for assessing the abundance and integrity of the protein regulated by the constitutive promoter. We selected the FLAG tag (DYKDDDDK) for purification because it had been used in conjunction with α-FLAG beads for purification and elution in previous TRAP protocols (Zanetti, Chang et al. 2005, Mustroph, Juntawong et al. 2009). We selected a Myc tag (EQKLISEEDL) for detection of the ribosomal protein because this tag works reliably in plant studies (Walter, Chaban et al. 2004). We also designed spacers in between GFP, tags, and RPL (supplemental figure 1). These sequences are intended to allow for interactions and re-folding of the linker without negatively impacting ribosome function or localization.

### Validation of Split GFP as an Efficient Linker

Our first experiments were designed to (1) compare the efficiency of split GFP reconstitution when expressed as different split fragments and with different positions of tags/spacers; (2) define the timeframe in which assembly of split GFP occurs; (3) observe whether the resultant protein complex localizes to the cytosol. These experiments employed transient expression via agroinfiltration in *Nicotiana benthamiana* (Voinnet, Rivas et al. 2003), followed by confocal microscopy for detection of GFP activity and western blotting to detect the FLAG and Myc tags.

Eight plasmid constructs were created in which every gene was placed under the control of the constitutive Cauliflower Mosaic Virus 35S promoter (*CaMV35S*) (Kay, Chan et al. 1987) (Figure 1c). The Arabidopsis RPL18b ribosomal gene was selected because it has been used successfully in previous TRAP protocols (Zanetti, Chang et al. 2005). Two different splits of the *GFP* open reading frame (Cabantous, Nguyen et al. 2013) were tested: a fragment encoding ß-sheets 1 through 9 (hereafter called *GFP1_9*) for refolding with a second fragment encoding ß-sheets 10 and 11 (*GFP10_11*); and a different split encoding ß-sheets 1 through 10 (*GFP1_10*), along with the corresponding 11^th^ ß-sheet (*GFP11*) (Figure 1c). Due to concerns that a large protein fused to the ribosomal protein could interfere with ribosome assembly or function, *RPL18b* was fused to the smaller fragment of each GFP split. GFP fragments were fused to the N-terminus of RPL18b, because the purification tag (Mustroph, Zanetti et al. 2009) and the GFP (Bailey-Serres personal communication) placed at the N-terminus of the RPL were shown to have no effect on ribosome assembly, function and purification. Two assemblies for each GFP fragment were created, positioning the Myc epitope either at the N-terminus of the GFP fragment-RPL fusion protein, or between the GFP fragment and the RPL (Figure 1c). For the second gene of the system, the FLAG tag was fused to the N-terminus of GFP1_9 and GFP1_10 to position the tag for immunoprecipitation. Two additional assemblies, *FLAG-GFP-Myc-RPL* and *FLAG-GFP* were constructed for use as positive controls.

Each pair of split GFP plasmids were co-infiltrated into leaves of *Nicotiana benthamiana*, and the timing, intensity, and subcellular localization of GFP activity in epidermal cells was assayed by microscopy. At 3 days post infiltration (dpi), fluorescence was detectable in all samples (Figure 2). Fluorescence increased in all samples over time. The GFP1_9/10_11 combination fluoresced stronger than the GFP1_10/11 combination. The position of the Myc tag seemed to affect fluorescence from the GFP1_10/11 assemblies: very little fluorescence was emitted from the samples expressing GFP1_10 + GFP11-Myc-RPL, compared to GFP1_10 + Myc-GFP11-RPL18b. The placement of the Myc tag in this configuration could interfere either with GFP re-folding, ribosome function, or ribosomal protein interactions, leading to reduced fluorescence. Both splits exhibited nucleo-cytoplasmic localization similar to the non-split FLAG-GFP-Myc-RPL18b control (Figure 2). These results indicate that the split GFP assembles quickly and fluoresces brightly without affecting RPL18b localization. The combination of FLAG-GFP1_10 + Myc-GFP11 appeared to be the best configuration for the system, because it fluoresced stronger than the other combinations.

**Figure 2.**
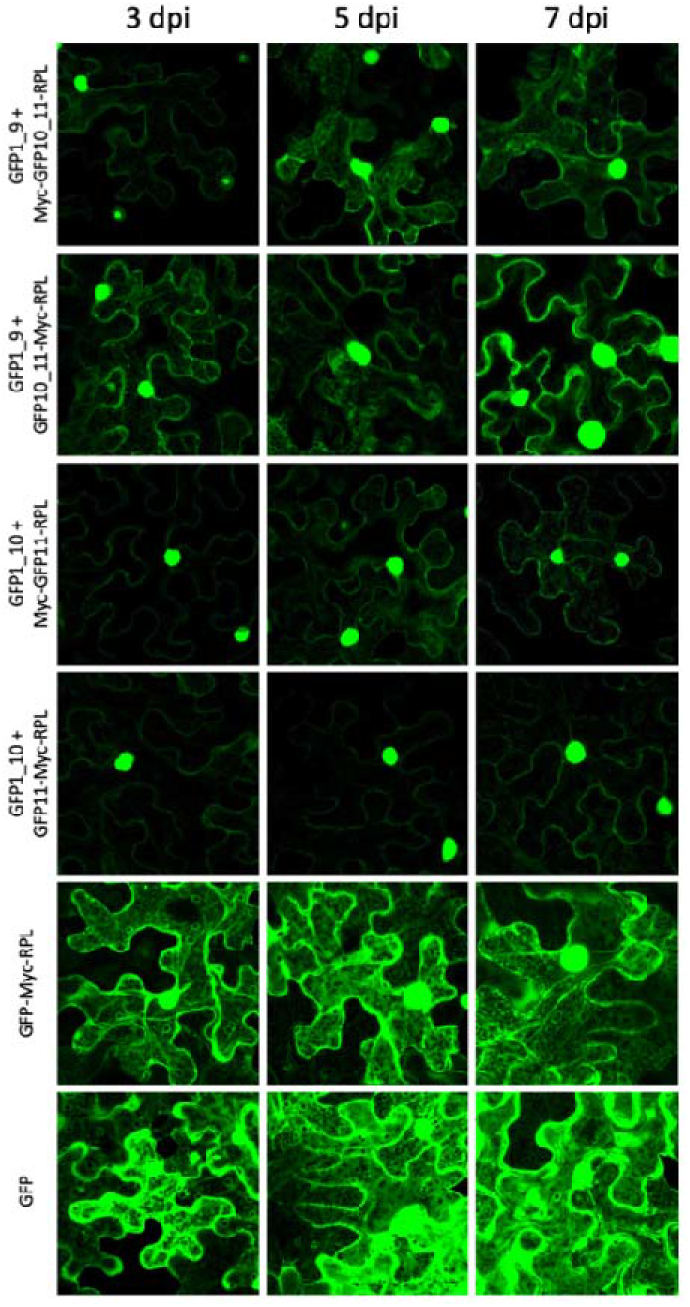
Comparison of fluorescence from GFP protein splits and different tag positions. Confocal microscopy was used to assay GFP activity and subcellular localization in *N. benthamiana* leaves transiently transformed through infiltration of *Agrobacterium* containing the genes of interest in T-DNA vectors. Images were recorded as z-stacks of multiple planes. “dpi” indicates number of days post infiltration with *Agrobacterium*; labels correspond to the constructs that are illustrated in Figure 1c.

### Validation of Assemblies for Two-Gene TRAP: Protein Expression and Purification

Our next set of experiments tested (1) whether the association between split fragments of GFP was strong enough to allow for recovery of the tagged ribosomes via immunoprecipitation; (2) whether ribosome yield was affected by the location of the GFP split or the position of the Myc tag. Infiltrated *N. benthamiana* leaves were harvested at 3 and 7 dpi, and three different samples from each time point were collected to track protein abundance at different steps in the immunoprecipitation procedure: The first sample was collected from <u>c</u>larified <u>l</u>eaf <u>e</u>xtract (CLE), comprised of the soluble fraction from leaf tissue homogenized in detergent buffer and clarified via filtration and centrifugation (Figure 3a). The second sample was collected from the resuspension of the pellet recovered after ultracentrifugation through a sucrose cushion (<u>S</u>ucrose <u>U</u>ltracentrifugation <u>P</u>ellet, SUP); only dense macromolecular complexes such as ribosomes and polyribosomes can penetrate the sucrose cushion (Figure 3b). The third sample was collected from molecules <u>i</u>mmuno<u>p</u>recipitated (IP) from the resuspended SUP with beads coated with anti-FLAG antibodies (Figure 3c). These samples were assayed via western blotting (Figure 4). An α-FLAG antibody was used to detect fusions with GFP1_9 and GFP1_10, as well as the full-length FLAG-GFP-Myc-RPL and FLAG-GFP controls. The α-Myc antibody was used to detect GFP10_11-RPL and GFP11-RPL, with each variant of Myc tag position.

**Figure 3.**
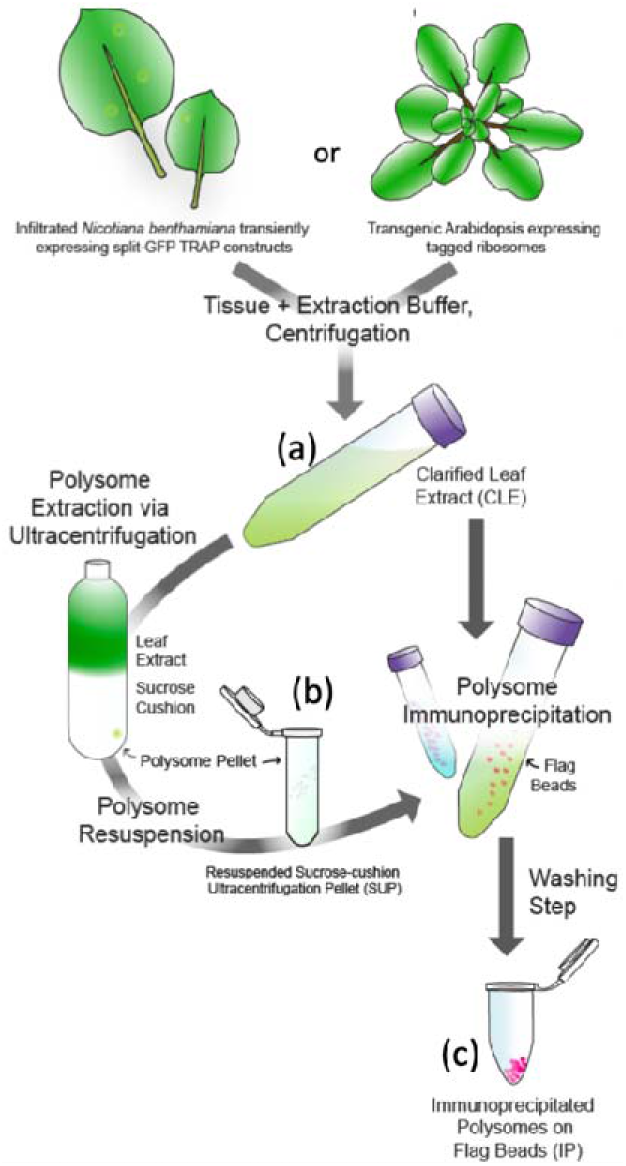
Schematic of the TRAP workflow. (a) Tissue from transiently transformed *N. benthamiana* or stably transformed Arabidopsis was collected, flash frozen, and homogenized with extraction buffer. Clarification of the homogenate via centrifugation yielded the clarified leaf extract (CLE), containing all soluble proteins. (b) Polysome concentration was achieved by ultracentrifugation of CLE over a sucrose cushion followed by resuspension, yielding the sucrose-cushion ultracentrifugation pellet (SUP) containing all protein complexes with sufficient density to penetrate the cushion. (c) Immunopurification of tagged polyribosomes from either CLE or SUP employed FLAG beads to yield immunoprecipitated polysomes (IP). Note that step (b) was omitted in some experiments (see text for details), in which the CLE was directly processed with Flag beads.

**Figure 4.**
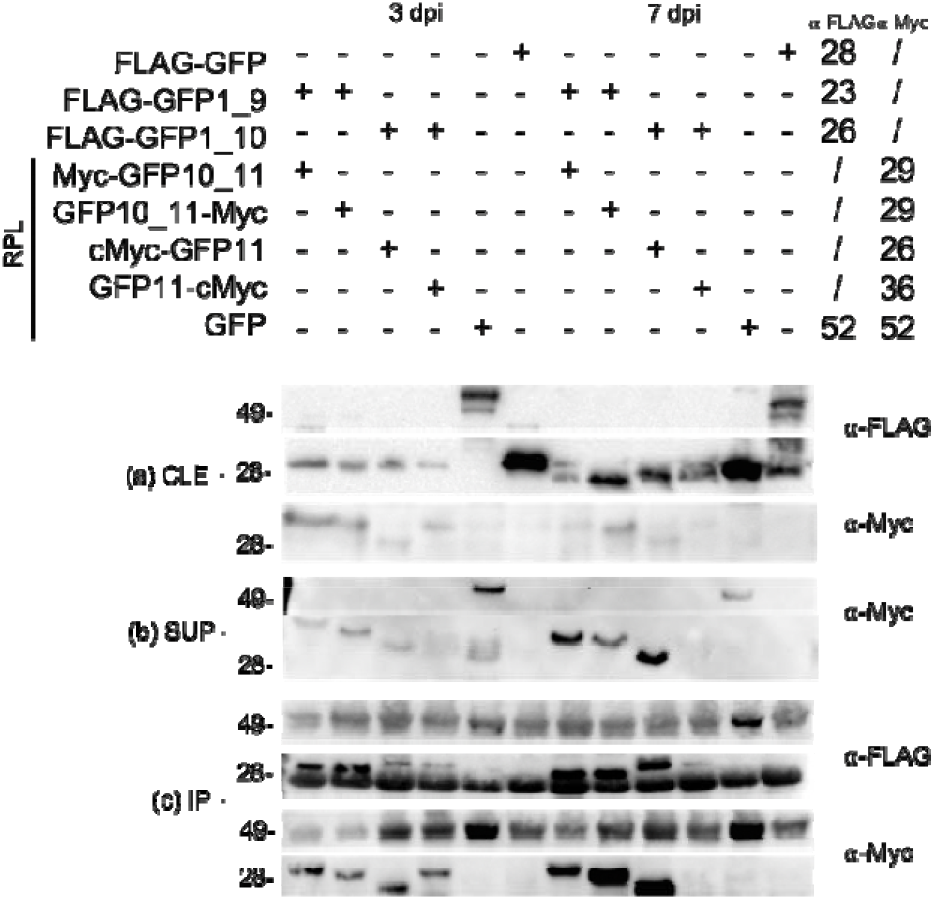
Western blot analysis of TRAP proteins from transient expression in *N. benthamiana*. (a) Samples from clarified leaf extract (CLE) at 3 and 7 dpi were probed with ⍰-FLAG to detect GFP1_9 or 1_10 fragments or ⍰-Myc to detect GFP10_11-RPL or GFP11-RPL fragments. (b) ultracentrifugation pellet (SUP) samples, probed with ⍰-Myc. No proteins were detected on this blot with ⍰-FLAG. (c) immunoprecipitated polysome (IP) samples probed with ⍰-FLAG or ⍰-Myc.The omnipresent bands at ~50 kda and ~25 kDa correspond to the ⍰-FLAG antibody from the beads, detected by the anti-mouse antibody. Top diagram: the numbers of the right correspond to the expected size of the fragments in kDa. Blots below: the number on the left of the blots correspond to the size of the molecular weight marker (kDa).

We expected to detect bands from every transiently expressed protein at both time-points from the CLE samples when probing with α-FLAG and α-Myc, because the CLE contains all proteins expressed in the leaf. Accordingly, proteins from every transgene were evident in samples from both 3 and 7 dpi (Figure 4a). As a control, we collected and assayed tissue infiltrated with only FLAG-GFP1_9 or only FLAG-GFP1_10. These proteins were detected with α-FLAG in CLE samples but not in the SUP or IP samples, as expected (not shown).

Only proteins from macromolecular complexes dense enough to penetrate the sucrose cushion should be detected in samples taken from the SUP or the IP. Transiently expressed proteins should be detected in these fractions only if they were incorporated into ribosomes at the time of tissue collection. Accordingly, the full-length Flag-GFP-Myc-RPL fusion protein control was detected with α-Myc in the SUP (Figure 4b) and IP (Figure 4c), while the Flag-GFP protein was not detected in either sample.

Each split GFP fusion protein should be detectable in SUP and IP only if it assembled with its cognate split, was incorporated into ribosomes, and if the association between the GFP fragments was sufficiently strong to withstand the sucrose centrifugation and subsequent immunoprecipitation with α-FLAG. Accordingly, all Myc-tagged GFP fragments were detectable with α-Myc in the SUP (Figure 4b) and IP (Figure 4c). Surprisingly, no Flag-tagged proteins were detectable in the SUP (not shown), perhaps due to the detergent-heavy nature of the sample preventing the monoclonal anti-FLAG antibody to bind the FLAG tag. However, the FLAG-tagged GFP fragments were detected with α-FLAG in the IP from the SUP, demonstrating that these proteins were indeed present in the SUP (Figure 4c). In both the IP and SUP, the α-Myc blots showed low abundance of the assemblies with GFP11-Myc-RPL (Figure 4b, c), possibly due to low expression of the proteins, consistent with the low fluorescence detected from this split (Figure 2).

These results demonstrate that while all tagged proteins were present in the CLE, only tagged proteins designed to incorporate into or link with ribosomes were present in the SUP and IP samples, implying their incorporation into ribosomes. As expected, free GFP was excluded from SUP and IP samples, due to its inability to penetrate the sucrose cushion. These data indicate that transiently expressed fusion proteins from our split linker TRAP system can assemble, are incorporated into ribosomes and can be immunoprecipitated.

### Validation of Assemblies for TRAP-RNAseq: RNA Extraction and Analysis

Having established that tagged ribosomal proteins could be immunoprecipitated from homogenized leaf tissue, we tested whether RNA could be isolated from these samples with quality and yield sufficient for downstream analyses. RNA from aliquots of the fractions used in western blotting from the CLE, SUP and IP was extracted and quantitated by spectrophotometry. RNA quality was assessed by both agarose gel electrophoresis and chip-based capillary electrophoreses. A greater amount of RNA was extracted from SUP samples compared to IP samples (Supplemental Table 1; 2-7.9 μg compared to 0.1-1.4 μg/sample). No RNA was detected from the IP of the free GFP samples, consistent with no detection of protein in western blotting (Figure 4c). The following RNA electrophoresis profiles were expected for CLE and SUP samples: 25S, 18S cytosolic rRNAs, and the 23S and 16S plastidial rRNAs, with the SUP samples enriched in the cytosolic rRNAs compared to the CLE samples. The IP samples were expected to contain only 25S, 18S cytosolic rRNAs, with little or no plastidial rRNAs because the corresponding ribosomes are not tagged. All samples displayed the expected patterns (Figure 5a). IP samples from GFP1_9 + Myc-GFP10_11-RPL, GFP1_9 + GFP10_11-Myc-RPL, as well as the GFP-Myc-RPL positive control exhibited higher RNA yields compared to GFP1_10 + GFP11-RPL splits, consistent with results from GFP imaging (Figure 2) and western blotting (Figure 4). RNA samples from the higher-yielding splits were selected for an additional validation step on an Agilent Bioanalyzer using the 7 dpi SUP and the 3 and 7 dpi IP samples (Figure 5b). All IP samples scored an RNA Integrity Number (RIN) of 8 or above indicating very good quality for sequencing. Each sample had a total RNA yield of 1 μg/sample or greater (Figure 5b), surpassing the minimum amount needed for most sequencing applications.

**Figure 5:**
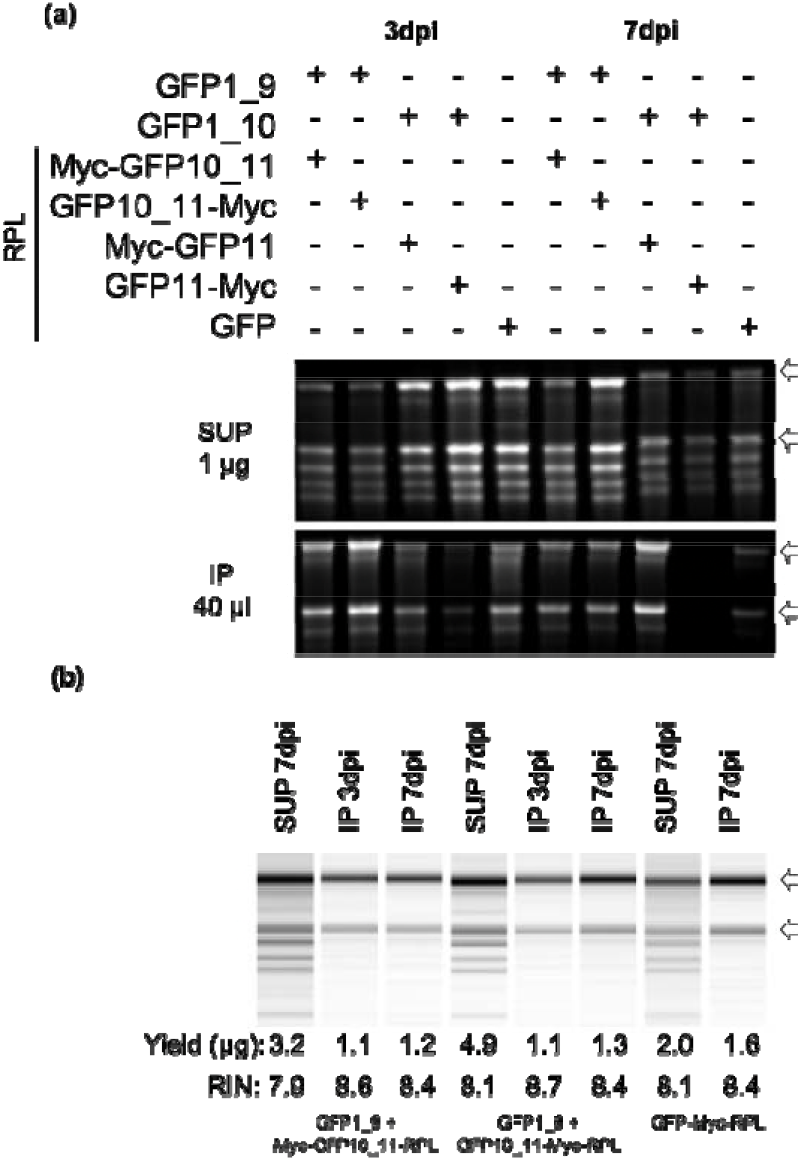
RNA analysis of SUP and IP RNA samples. RNA gel electrophoresis of samples from sucrose ultracentrifugation pellet (SUP) and immunoprecipitation (IP) fractions at 3 and 7 dpi. (a) SUP samples. One μg of RNA from each sample was dried, resuspended with RNA loading buffer, and resolved on a 1.2% agarose gel. (b) IP samples. The total amounts of RNA extracted were dried, resuspended, and loaded. (c) Capillary electrophoresis analysis displays banding patterns from selected SUP and IP samples, with yield and RNA Integrity Number (RIN) scores from each sample. Open arrows points towards the 25S and 18S rRNAs.

The data from fluorescence, western blotting, and RNA analyses indicated that GFP1_9 + GFP10_11 assembly pair performed better than GFP1_10 + GFP11 assembly pair, possibly resulting from issues with GFP refolding or protein accumulation. The position of the Myc tag had little influence on the RNA quality and yield, although FLAG-GFP1_9 + Myc-GFP10_11-RPL led to slightly higher fluorescence than the FLAG-GFP1_9 + GFP10_11-Myc-RPL combination (Figure 2). We therefore selected FLAG-GFP1_9 + Myc-GFP10_11-RPL for application of the split GFP TRAP technology to Arabidopsis.

### Selection and Validation of Arabidopsis Promoters

The split GFP TRAP system was tested to study the interactions between Arabidopsis and the oomycete pathogen *Hyaloperonospora arabidopsidis* (*Hpa*), in order to determine the translatome from plant cells that contain pathogen feeding structures called haustoria, present primarily in mesophyll cells (Herlihy, Nora et al. 2019). Therefore, we searched the literature for promoters that would provide the desired cell specificity for the two gene fusions: a constitutive, mesophyll-specific promoter for the expression of Myc-GFP10_11-RPL, and an Hpa-responsive promoter to drive the expression of FLAG-GFP1_9 specifically in haustoriated cells, to specifically tag ribosomes in haustoriated cells. Two promoters fulfilled these criteria: the *PLASTOCYANIN 2* (*PETE2*, AT1G20340) promoter, as a 1790 bp-long DNA sequence that was previously shown to be predominantly active in the mesophyll (Vorst, van Dam et al. 1993); and the promoter for Arabidopsis *<u>D</u>owny <u>M</u>ildew <u>R</u>esistant 6* (*DMR6*, ATG5G24530), which encodes a defense-associated gene required for susceptibility to *Hpa*, and was previously reported to be active almost exclusively in haustoriated cells (Van Damme, Huibers et al. 2008).

To validate the localization of activity from these promoters under our experimental conditions, the *PETE2p* and *DMR6p* promoters were fused to the *E. coli* ß-Glucuronidase (*GUS*) gene and introduced into Arabidopsis. 3-week-old *PETE2p-GUS* transgenic plants exhibited uniform GUS activity in the mesophyll, with an absence of activity in all major veins (Figure 6a). *DMR6p-GUS* transgenic plants exhibited GUS activity from 3 days post-infection (dpi) with *Hpa* onwards in cotyledons of seedlings (Figure 6b). However, GUS staining was not restricted to haustoriated cells in seedlings. Contrastingly, in infected true leaves of 3-week old plants, GUS staining was almost exclusively confined to haustoriated cells (Figure 6c; supplemental Figure 2). We consequently analyzed true leaves from adult plants rather than cotyledons from seedlings.

**Figure 6:**
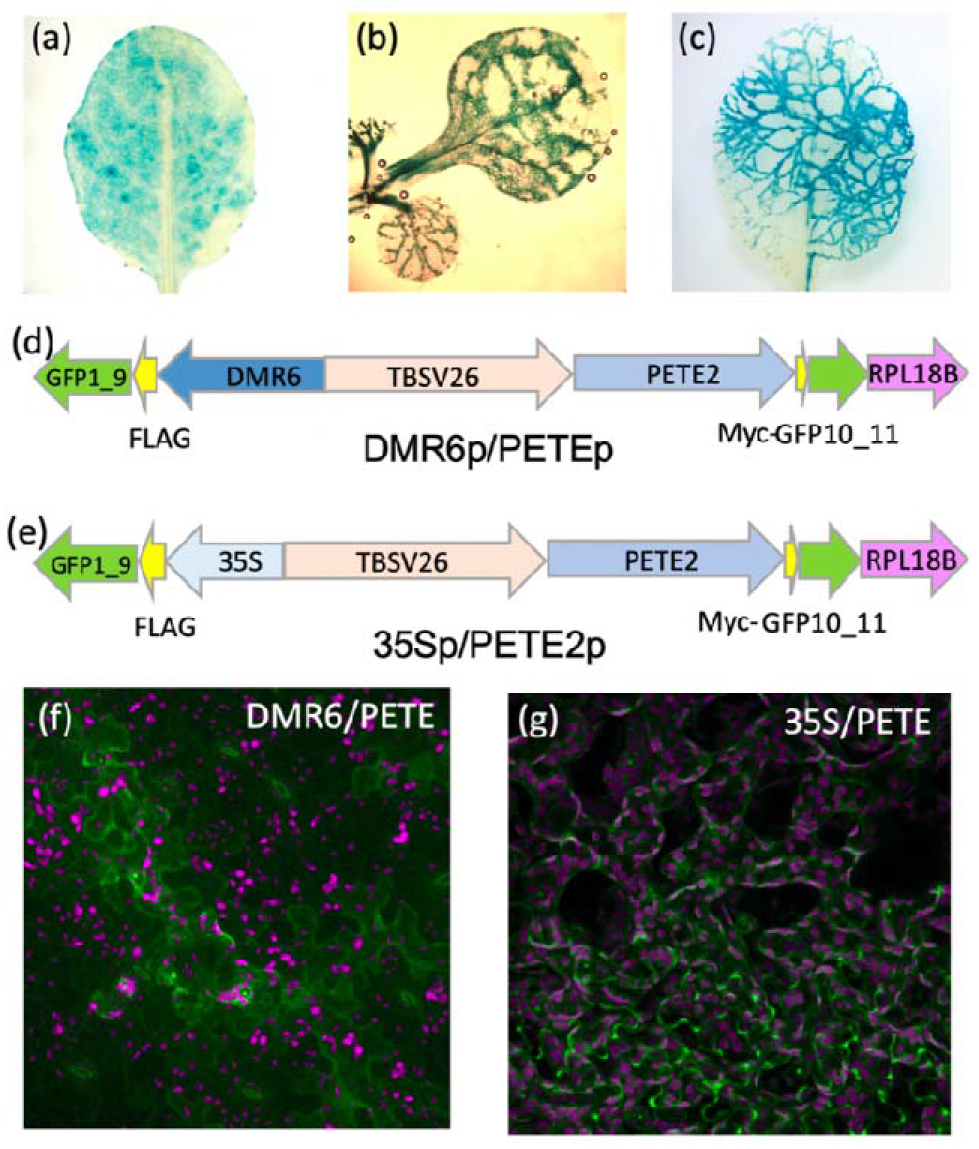
Characterization of *PETE* and *DMR6* promoter activity. (a-c) Histochemical staining for β-Glucuronidase (GUS) activity in soil-grown, transgenic Arabidopsis expressing (a) *PETEp-GUS* in a true leaf from a three-week-old plant. (b) *DMR6p-GUS* in *Hpa*-infected cotyledons from a 16-day-old seedling at 6 dpi. (c) *DMR6p-GUS* in *Hpa*-infected true leaves from a three-week-old plant at 6 dpi. (d, e) Maps of plasmid for (d) *DMR6/PETE* and (e) *35S/PETE*. (f, g) Confocal microscopy images of infected three-week old transgenic Arabidopsis expressing (f) *DMR6/PETE* at 6 dpi (infected with Hpa) and (g) *35S/PETE* (uninfected).

### Trials of the split GFP TRAP system with Arabidopsis infected by *Hpa*

For the use of TRAP of haustoriated cells in Arabidopsis, a single plasmid was constructed with two assemblies: PETE2p-Myc-GFP10_11-RPL and DMR6p-FLAG-GFP1_9, separated by the TBSV26 insulator to prevent promoter interference (Hily, Singer et al. 2009) (Figure 6d). This plasmid was named *“DMR6p/PETEp”*. A second plasmid, named *“35Sp/PETEp”*, was constructed to collect ribosomes from all mesophyll cells, regardless of pathogen presence or absence (Figure 6e). This plasmid was identical to *DMR6p/PETEp* except that the *DMR6p* promoter was replaced by 35Sp. In infected *DMR6p/PETEp* plants, GFP activity was detected in haustoriated cells, demonstrating that the DMR6p promoter drives the expression of *FLAG-GFP1_9* in the presence of the pathogen, and that GFP1_9 refolds with *PETE2p*-driven Myc-GFP10_11-RPL (Figure 6f). No fluorescence was detected in uninfected *DMR6p/PETEp* lines (not shown). As expected, the *35Sp/PETEp* plants showed strong fluorescence in all mesophyll cells (Figure 6g).

Tissue from *Hpa*-infected and mock-infected *DMR6p/PETEp* and *35Sp/PETEp* lines was collected for TRAP. The infection experiment was repeated 15 times so that the most similar replicates could be chosen for analysis, thereby reducing the noise from biological variability in the Arabidopsis-Hpa interaction. Quality control assays were run on each of the 15 replicates, and 4-5 replicates for each condition where selected for TRAP and RNA extraction (Supplemental Figure 3). Two RNA samples, respectively comprising the transcriptome and translatome, were taken from each replicate: one from the CLE and one from the IP. The TRAP RNA extraction differed slightly from the protocol used for samples from infiltrated *Nicotiana benthamiana*: we did not include the sucrose cushion step, and the CLE samples were incubated twice with FLAG beads to increase RNA yields. We observed a higher yield and the typical ribosomal banding pattern from the total RNA sample, and a lower yield and the expected ribosomal banding pattern from TRAP RNA sample (9-15 vs. 1-10.9 μg/sample respectively; Figure 7; supplemental figure 4; supplemental table 2). RNA yields from the *DMR6p/PETEp* lines ranged from 1.1 to 1.3 μg/sample. This relatively low yield was anticipated due to the limited number of haustoriated cells in the leaves. As expected, no TRAP RNA was recovered from uninfected *DMR6p/PETEp*, because the *DMR6* promoter is inactive under those conditions. The quality and quantity of RNA from all other samples were sufficient for RNA sequencing. Analyses of these samples will be described in a forthcoming manuscript.

**Figure 7.**
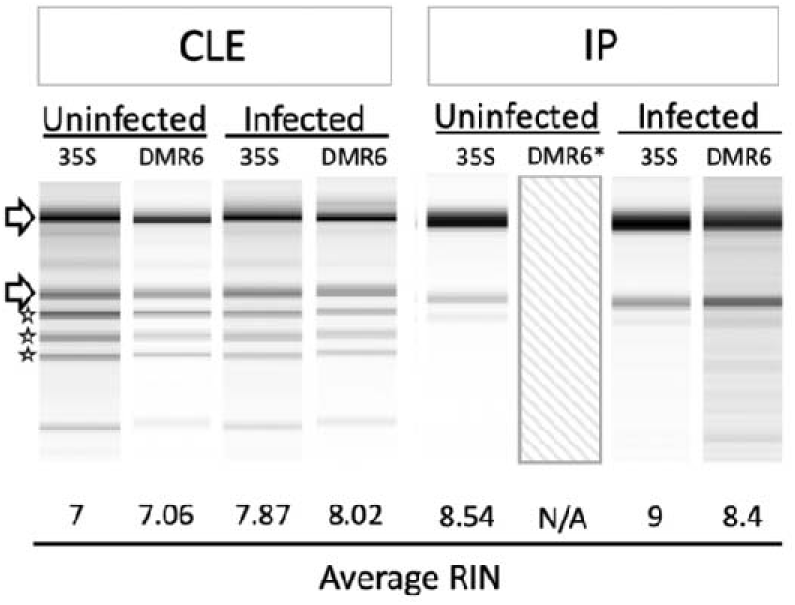
Capillary electrophoresis of RNA from transgenic Arabidopsis. Banding patterns of RNA from clarified leaf extract (CLE) and immunoprecipitated polysome (IP) samples from infected and uninfected plants. “35S” denotes the *35S/PETE* line, while “DMR6” denotes the *DRM6/PETE* line. Average RIN is calculated from five samples, and a representative image was chosen for each condition. *No image or RIN is present for uninfected IP samples from the DMR6 line because no tagged ribosomes were recovered from the immunopurification of this sample. Arrows point towards the 25S and 18S rRNAs, while the stars indicate the organelle rRNAs.

## Conclusions

We have designed a new system applicable to any cell-specific or stimulus-specific TRAP in plants, based on a split GFP linker. We applied this system to Arabidopsis-*Hpa* interactions, using promoters that enable specific enrichment of ribosomes from haustoriated cells from a heterogeneous, whole-leaf sample of haustoriated and non-haustoriated cells. To demonstrate the viability of this design, we optimized and validated several important features: (1) Split superfolder GFP can re-fold efficiently in our system, providing an approach for adding purification tags to pre-existing ribosomes for efficient recovery via immunoprecipitation. GFP florescence was also validated as a visual marker for expression and assembly of the fragments of the split GFP. (2) The best split of the GFP gene was between sheets 9 and 10, and the best configuration of linkers and tags was FLAG-GFP1_9 + GFP10_11-Myc-RPL. (3) Association of the split GFP fragments is strong enough to withstand centrifugation through a sucrose cushion, demonstrating that the GFP10_11-Myc-RPL fusion protein will not dissociate from the FLAG-GFP1_9 protein during TRAP. (4) The amount and quality of the RNA extracted from immunoprecipitated ribosomes are sufficient for qPCR or RNA sequencing. We expect that these findings should be broadly applicable to any application of the split GFP system for TRAP in plants and potentially other organisms.

With respect to our goal of obtaining the translatome of haustoriated cells, we established that (1) The *DMR6* promoter is activated specifically in haustoriated plant cells only in true leaves, not in cotyledons. (2) The *DMR6p/PETEp* and *35Sp/PETEp* systems performed well in transgenic Arabidopsis infected with Hpa: immunoprecipitation of ribosomes from the CLE allowed for purification/concentration of RNA in amounts and quality sufficient for RNA sequencing. Our subsequent analyses of the translatomes from haustoriated cells, to be published elsewhere, confirmed that this system provides new insights into plant-oomycete interactions, and we expect that this approach could be applied productively to many different plant-pathogen interactions, provided that a specific pathogen-inducible promoter is available for the interaction in question.

Existing TRAP technology provides a powerful tool for researchers to interrogate the translatome. Previous applications of TRAP have employed a single transgene, driven by a strong, constitutive promoter, or less commonly, by a promoter that is active in specific cell types or in response to specific conditions to produce a tagged ribosomal protein that enables TRAP (Urquidi Camacho, Lokdarshi et al. 2020). Our two-gene system enables cell-type or condition-specific attachment of a tag to an existing pool of ribosomes through refolding of a split GFP linker. This feature will increase the sensitivity of TRAP in situations where a tagged ribosomal protein from a single gene is unlikely to be incorporated into ribosomes with sufficient efficiency for robust yields (*e.g.* when ribosome protein turnover is slow or suppressed, or when the biological material is to be analyzed shortly after induction). Moreover, the two-gene system can provide an additional level of specificity through customization of two promoters instead of just one. Altogether, we anticipate that this system will be applicable to diverse biological contexts in which TRAP is applied to profile a specific target cell population from a complex assemblage.

## Experimental Procedures

### *Arabidopsis thaliana* and *Nicotiana benthamiana* Growth and Transformation

*Arabidopsis thaliana* and *Nicotiana benthamiana* were grown under 120 μE/m^2^/s, 22°C, 16 h light /8 h dark on soil (Sunshine Mix #1, Sungro Horticulture) and were watered from below with 300 mg/l Miracle-Gro Fertilizer (24-8-16 NKP, Scotts). Arabidopsis Ecotype Columbia 0 (Col-0, CS70000) was used for all experiments. Stably transformed Arabidopsis lines were generated by floral dipping using *Agrobacterium tumefaciens* GV3101 (pMP90) (Clough Steven and Bent Andrew 2008). For transient expression in *N. benthamiana*, leaves of 5-week old plants were infiltrated with a suspension of *Agrobacterium tumefaciens* GV3101 (pMP90) carrying the constructs or p19 as in Batoko et al. 2000 (Batoko, Zheng et al. 2000, Yu and Pilot 2014) with the following modifications: the bacteria were grown overnight in LB medium supplemented with appropriate antibiotics, washed twice in 10 mM MgCl_2_, 100 μM acetosyringone, and diluted to final OD_600_ of 0.05 in the same solution.

### *Hpa* Propagation and Infection

*Hpa* isolate Noco2 was used for all experiments, as described in (Reignault, Frost et al. 1996, McDowell, Hoff et al. 2011). Sporangiospore suspensions of 5 × 10^4^ spores/ml were prepared in water from sporulating plants and sprayed on seedlings or 3-week-old Arabidopsis plants. Inoculated plants were covered overnight, and then uncovered and kept under short-day (8h, 22°C/16h, 20°C day/night) conditions. Control plants were sprayed with water alone. Infected plant material for TRAP was harvested into liquid nitrogen at 6 dpi, prior to sporulation.

### GUS and Trypan Blue Staining

GUS staining was performed with fresh leaf tissue. Leaves were fixed under vacuum in sodium phosphate buffer (50 mM, pH 7.2) containing 0.5% Triton and 1.5% formaldehyde for 45 min. Fixed leaves were washed three times for 5 min in phosphate buffer, 0.5% Triton with 500 μM each of potassium ferrocyanide and potassium ferricyanide and infiltrated with wash buffer as above containing 1 mM X-Gluc (GoldBio). Samples were incubated at 37°C in the dark until staining became visible and were cleared with ethanol for imaging. Trypan blue staining of Hpa hyphae, haustoria, and spores was performed as described in (McDowell, Hoff et al. 2011) with the following modification: leaves were incubated for 90 seconds at 90°C, incubated at room temperature for 5 min, and de-stained with chloral hydrate.

### Fluorescence Microscopy

Prior to imaging, leaves were infiltrated with water via syringe or vacuum and placed on slides with abaxial side facing the slide cover. Imaging was performed on a Zeiss LSM 880 confocal laser scanning microscope, with Zen Black software.

### Western Blotting

Aliquots of extracts takes at different stages of the TRAP procedure were prepared with 1 M DTT and 4X NuPAGE LDS sample buffer, denatured at 90°C for 10 min, and analyzed by SDS-PAGE (4-12% polyacrylamide NuPAGE MES gels, Life Technologies). Proteins were transferred to nitrocellulose membrane (GE Healthcare) by wet electroblotting. The filter was treated with Ponceau red stain and blocked in OneBlock (Genesee Scientific). Proteins were detected using primary antibodies α-FLAG (monoclonal α-FLAG M2, Sigma, 1:1000) or α-Myc (Clone A-14, Santa Cruz; 1:1000) and secondary antibody Horse Radish Peroxidase (HRP)-α-mouse (ThermoFisher Scientific, 1:10,000) in OneBlock buffer. HRP activity was detected using ECL-Plus western blotting detection system (GE Healthcare), with luminescence captured by a ChemiDoc CCD camera imaging system (Biorad).

### Multiple Fragment Cloning using In-Fusion

Primers were designed for each fragment with 10-20 bp overlaps corresponding to the fragments that would be adjacent to the amplified fragment in each corresponding clone. Fragments were amplified from gBlocks Gene Fragments (Integrated DNA Technologies), Col-0 genomic DNA or existing fragments by PCR with the KOD Hotstart DNA polymerase (Toyobo). Amplified fragments were gel purified and used for In-Fusion Multiple Fragment Cloning (Clontech) according to manufacturer’s directions. Clones were verified by sequencing.

### Translating Ribosome Affinity Purification

TRAP was performed according to Mustroph et al. (Mustroph, Juntawong et al. 2009) with the following modifications: For *N. benthamiana* samples, 8 ml of extraction buffer was added to 1.5 g of tissue ground in liquid N_2_. The resulting leaf extract was clarified twice via centrifugation, with a miracloth filtering step between centrifugations. 6 ml of clarified leaf extract was loaded on top of a 1.7 M sucrose cushion containing detergents (DOC, PTE, and Detergent Mix) in the same concentrations as in the extraction buffer. Following centrifugation, the pellet was resuspended in 500 μl of extraction buffer by agitation on a spinning wheel at 4°C overnight, followed by removal of insoluble debris by centrifugation for 1 min, 8200×g at 4°C. The resuspended polysomes were added to extraction buffer to a final volume of 5 ml with the addition of 100 μl washed FLAG beads (Sigma). Polysomes were incubated with the beads for 2 h with gentle rocking at 4°C. The beads were washed as follows: one wash with 5 ml extraction buffer, 4 washes with 5 ml wash buffer with 5 min incubation periods between washes. For Arabidopsis samples, clarified leaf extract was added directly to previously washed FLAG beads, and followed the same incubation and wash protocol as above. To account for a lower number of tagged ribosomes, clarified leaf extract samples from the Arabidopsis *DMR6p/PETEp* line were saved and used for a second consecutive incubation with an additional 100 μl of washed FLAG beads.

### RNA Extraction and Quantification

RNA was extracted from Arabidopsis tissue, clarified leaf extract, resuspended polysomes, or FLAG-beads using either 1 ml of TRI REAGENT (Sigma) or the ISOLATE II RNA Plant kit (Bioline, Meridian Life Science). RNA integrity was confirmed by either agarose gel electrophoresis or by capillary electrophoresis in an Agilent 2100 Bioanalyzer according to manufacturer’s instructions. cDNA was synthesized from 1.5 μg of RNA using the SuperScript IV Reverse Transcriptase kit (ThermoFisher Scientific) according to manufacturer’s instructions in a 10 μl reaction volume. Quantitative PCR was performed with 5 μl of the product (diluted 50 times in water), 5 μl of primer mix (1 μM each), and 10 μl of 2X PowerUp SYBR Green Master Mix (Applied Biosystems) on the Applied Biosystems 7500 Real-Time PCR System (2 min 50°C, 10 min 95°C, 40 cycles of [15 sec 95°C, 15 sec 55°C, 1 min 72°C]).

### Infected Plant Nucleic Acid Isolation and Pathogen Quantification via RT-PCR

Arabidopsis tissue infected with *Hpa* was harvested at 6 dpi and ground to a fine powder in liquid nitrogen. DNA was extracted using the BioSprint 96 DNA Plant kit (Qiagen) according to manufacturer’s specifications. The protocol from Anderson et al. (Anderson and McDowell 2015) was followed with modifications for the use of TaqMan polymerase, reagents, and probes instead of SYBR Green reagents (Applied Biosystems).

## Acknowledgments

This research was supported by funding from the National Science Foundation (IOS-1353366) for JMM and GP, and the Hatch Program of the National Institute of Food and Agriculture (VA-135908 for GP and VA-160106 for JMM) and the Virginia Agricultural Experiment Station. The authors have no conflict of interest to declare.

## Supplemental Tables

**Supplemental table 1:** Total yield (μg) of RNA extracted from SUP and IP *Nicotiana benthamiana* samples. For each construct combination at the SUP and IP experimental timepoints, at 3 and 7 dpi, RNA was extracted and quantified via capillary electrophoresis.

|  | Sample | 3 dpi | 7 dpi |
| --- | --- | --- | --- |
| SUP | GFP1_9 + Myc-GFP10_11-RPL | 4.19 | 3.29 |
|  | GFP1_9 + GFP10_11-Myc-RPL | 2.6 | 4.94 |
|  | GFP1_10 + Myc-GFP11-RPL | 3.03 | 3.41 |
|  | GFP1_10 + GFP11-Myc-RPL | 7.86 | 6.28 |
|  | GFP-Myc-RPL | 2.01 | 2.15 |
| IP | GFP1_9 + Myc-GFP10_11-RPL | 0.89 | 1 |
|  | GFP1_9 + GFP10_11-Myc-RPL | 0.91 | 1.04 |
|  | GFP1_10 + Myc-GFP11-RPL | 0.58 | 1.36 |
|  | GFP1_10 + GFP11-Myc-RPL | 0.33 | 0.08 |
|  | GFP-Myc-RPL | 0.68 | 1.29 |

**Supplemental table 2:** RNA yield (μg) and quality from Arabidopsis TRAP conditions. Average (Avg) total yield and standard deviation (Stdev) of total RNA and average RIN scores from 4-5 replicates of each line and infection status used in the Arabidopsis TRAP experiment. RNA was extracted by column purification and quality was measured via capillary electrophoresis.

| Condition | Avg Yield | Stdev | RIN |
| --- | --- | --- | --- |
| 35S/PETE IP- Mock | 9.4 | 1.63 | 8.5 |
| 35S/PETE IP- Infected | 6.5 | 0.7 | 9 |
| DMR6/PETE IP- Infected | 1.4 | 0.24 | 8.4 |
| 35S/PETE CLE- Mock | 12.3 | 1.59 | 7 |
| DMR6/PETE CLE- Mock | 12.6 | 2.5 | 7.9 |
| 35S/PETE CLE- Infected | 10.5 | 1.1 | 7.1 |
| DMR6/PETE CLE- Infected | 13 | 1.25 | 8 |

**Supplemental Figure 1.**
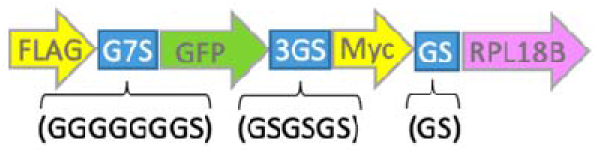
Illustration of amino acid spacers placed between components of TRAP assemblies. Seven glycine codons were placed between the FLAG tag and GFP (G7S), three repeats of Glycine-Serine were placed between GFP and the Myc tag (3GS), and one G-S repeat was placed between the Myc tag and RPL18B (GS).

**Supplemental Figure 2.**
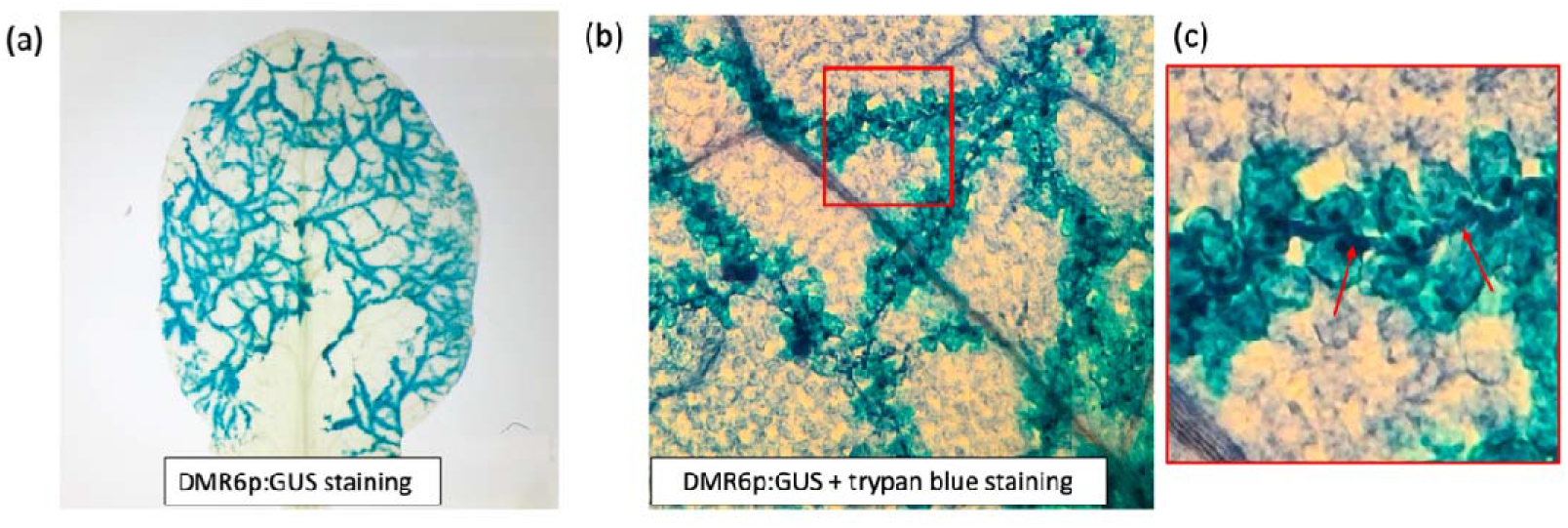
*DMR6p* activity in 3-week-old Arabidopsis leaves infected with *Hpa*. (a) Staining for β-Glucuronidase activity from plants expressing a *DMR6p-GUS* construct illustrates the pattern of *DMR6* promoter activity in 3-week-old true leaves at 6 dpi. (b) Dual staining for β-Glucuronidase activity and trypan blue illustrates *DMR6* promoter activity adjacent to *Hpa* hyphae and haustoria. (c) Enlargement of the segment in (b) bordered by red lines. Red arrows point to haustoria.

**Supplemental Figure 3.**
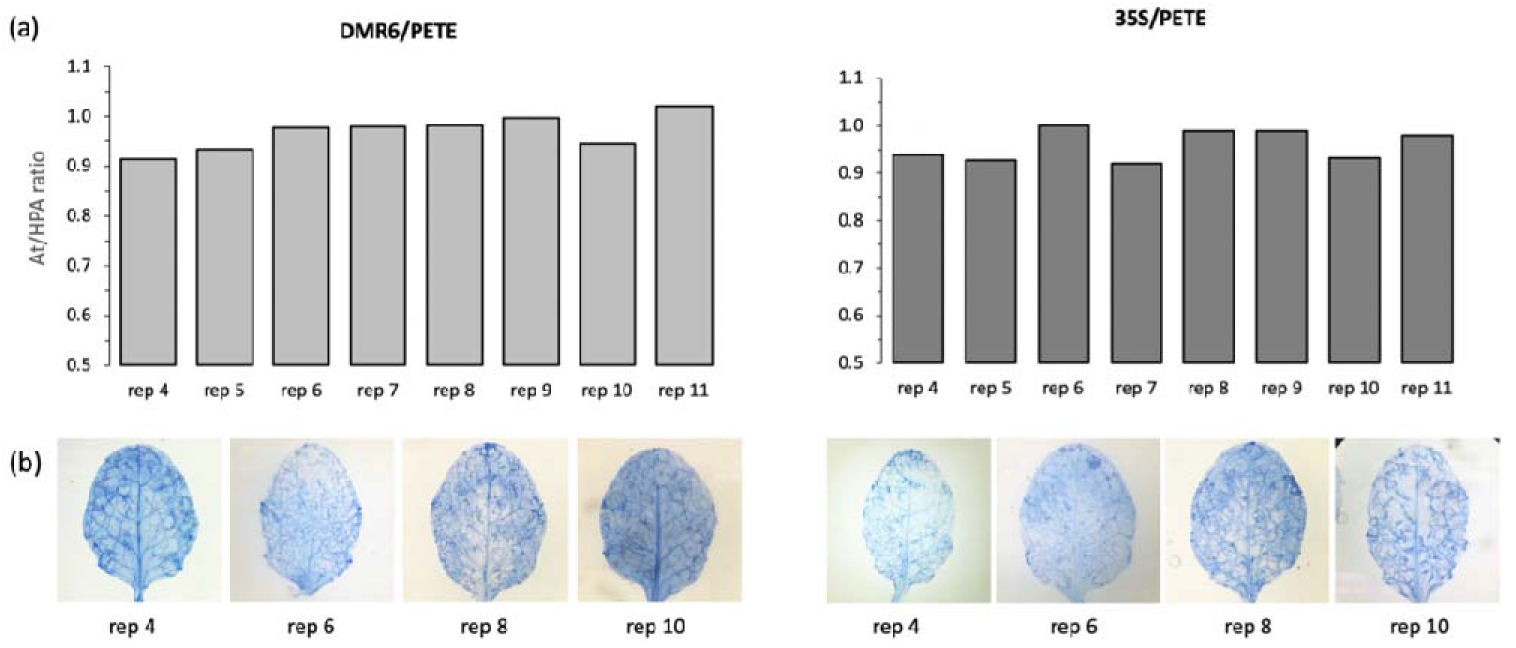
Pathogen biomass assays for replicate selection. (a) A quantitative PCR assay to determine vegetative growth of *Hpa* in plants from each infected replicate. Ratios were calculated from the cycle numbers for the *A. thaliana* (At) Actin2 amplicon and the *Hpa* actin amplicon. (b) Trypan blue staining illustrates *Hpa* hyphal growth in representative leaves from several replicates.

**Supplemental Figure 4.**
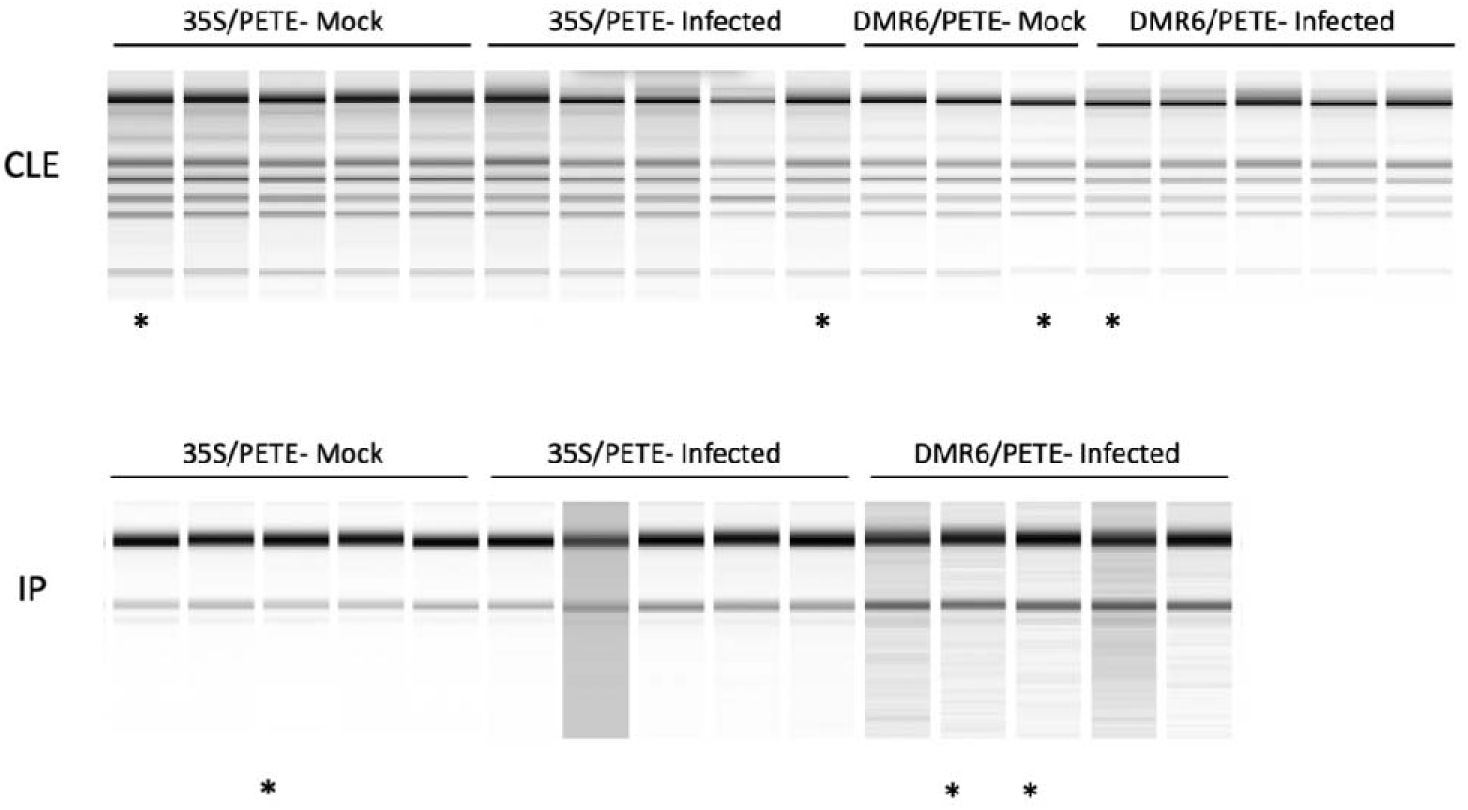
Capillary electrophoresis analysis of RNA from transgenic Arabidopsis. Banding patterns of RNA from CLE and IP samples in infected and mock-treated samples from capillary electrophoresis. The asterisks denote samples shown in Figure 7.

